# Reversible regulation of Cas12a activities by AcrVA5-mediated acetylation and CobB-mediated deacetylation

**DOI:** 10.1101/2021.11.16.468554

**Authors:** Xiaoman Kang, Lei Yin, Songkuan Zhuang, Tianshuai Hu, Zhile Wu, Guoping Zhao, Yijian Chen, Yong Xu, Jin Wang

## Abstract

The clustered regularly interspaced short palindromic repeat (CRISPR)/CRISPR associated (Cas) system protects bacteria and archaea from the invasion of foreign genetic elements. To cope with the host CRISPR systems, phages have evolved many strategies, including the anti-CRISPR (Acr) proteins, to inactivate the Cas nucleases. Recently, it has been reported that the type V-A Cas12a effector can be acetylated and inactivated by AcrVA5, which is a GNAT-family acetyltransferase. However, it is unclear whether the host has any coping strategies to reactivate the defense system. Here we show that the AcrVA5-acetylated Cas12a can be deacetylated by bacterial deacetylase CobB, reactivating Cas12a for both in vitro cleavage of target DNA sequences and in vivo protection of the host from invasion of foreign nucleic acids. Therefore, this study not only shows the reversible regulation of Cas12a activities by post-translational modification but also reveals CobB as a secondary safeguard to bacterial CRISPR defense systems. In addition, we demonstrate that AcrVA5 is a wide-spectrum acetyltransferase, acetylating a large number of target proteins besides Cas12a, and the AcrVA5-acetylated targets can also be deacetylated by CobB.

## Main Text

The clustered regularly interspaced short palindromic repeat (CRISPR)/CRISPR associated (Cas) system protects bacteria and archaea from mobile genetic elements (MGEs) such as bacteriophages.^1^ The Cas effector can be guided by an CRISPR RNA (crRNA) to target the invading nucleic acids by base-pairing and then cleaves the target to provide the host with immunity.^2, 3^ To cope with the immunity pressure imposed by the host CRISPR system, phages have evolved different kinds of anti-CRISPR (Acr) systems to inactivate the Cas nucleases and close the host immunity system in turn.^4^ The Acr proteins can directly interact with the Cas proteins to prevent the crRNA loading or target binding; alternatively, they possess enzymatic activities to cleave the crRNAs or post-translationally modify the Cas proteins.^4^ Among them, there exists a recently reported GNAT-family acetyltransferase, namely AcrVA5, which is encoded by the MGEs and specifically inhibits the type V-A Cas12a effector through acetyl modification of a key lysine site. The AcrVA5-inactivated Cas12a loses the ability to interact with the protospacer adjacent motif (PAM) site and fails in protecting the hosts by cleaving the invading MGEs.^5, 6^ Through inactivating the Cas12a system, MGEs containing the *acrVA5* gene can effectively shut down the bacterial immunity system and have the potential to widely spread among the Cas12a-harboring hosts. However, to our great surprise, the *acrVA5* gene exists only in three *Moraxella bovoculi* strains which have lost their deacetylases (**Table S1**). Therefore, it is reasonable to suspect that there may exist a competitive relationship between the acetyltransferase AcrVA5 and the widely distributed bacterial deacetylases,^7^ which may reactivate Cas12a by deacetylation to protect the hosts from the invasion of MGEs harboring the *acrVA5* cassette.

We first performed the AcrVA5-mediated *in vitro* acetylation experiment and showed that the AcrVA5-acetylated *Lachnospiraceae bacterium* (Lb) Cas12a lost both *cis*- and *trans*-cleavage activities towards double-stranded target DNA (dsDNA) (**Figure S1a** and **S1b**), which was consistent with the previous findings;^5, 6^ however, the AcrVA5-treatment showed no effect on LbCas12a *trans*-cleavage activities when triggered by target single-stranded DNA (ssDNA) (Figure S1c). As the PAM site is only necessary for Cas12a to recognize target dsDNA but not ssDNA,^8^ the above results further confirmed that AcrVA5-mediated acetylation prevented Cas12a from interacting with the PAM sequences in target dsDNA.^5^

Besides LbCas12a, we also analyzed Cas12a orthologs from *Francisella tularensis* subsp. novicida (FnCas12a) and *Acidaminococcus sp.* (AsCas12a) and demonstrated that AcrVA5 was able to inactivate both orthologs (**Figures S2** and **S3**). Noticeably, AcrVA5 was ever shown to be ineffective against AsCas12a in a previous study;^6^ however, we found that AsCas12a contained the conserved lysine residue (**Figure S3c**) and showed that the AcrVA5-mediated treatment led to the loss of both *cis*- and *trans*-cleavage activities of AsCas12a with target dsDNA.

Post-translational lysine acetylation plays an important role in diverse cellular processes in organisms from bacteria to human. In bacteria, the NAD^+^-dependent sirtuin-type CobB deacetylates a large number of proteins and regulates the global acetylation level.^7^ To test whether CobB can deacetylate the AcrVA5-treated Cas12a and reactivate its *cis*- and *trans*-cleavage activities, we then purified recombinant *E. coli* CobB and LbCas12a and performed the *in vitro* deacetylation assay. Based on the western blot results, Cas12a was successfully acetylated by AcrVA5 at the presence of acetyl-CoA, and the acetyl modification could be efficiently removed after being treated by CobB. Consistent with previous findings^7^, the CobB-mediated deacetylation stringently requires NAD^+^ as the cofactor (**Figure S4**). Accordingly, the AcrVA5-acetylated LbCas12a lost both *cis*- and *trans*-cleavage activities with target dsDNA but fully recovered both activities after being deacetylated by CobB (**Figure 1a** and **1b**). In addition, we tested several Cas12a orthologs such as FnCas12a and AsCas12a, and found CobB was able to reactivate both Cas12a orthologs (**Figures S5** and **S6**). It was worthy to mention that the successful reactivation of acetylated AsCas12a by CobB treatment once again proved AsCas12a as a target of AcrVA5 (**Figure S3**), while the reason for the distinct results between this work and the previous study^6^ was still unknown and could be an interesting question subject to further investigation. Based on above results, one may conclude that AcrVA5 and CobB reversibly regulate both the *cis*- and *trans*-cleavage activities of Cas12a by modulating its acetyl status *in vitro*.

**Fig. 1.**
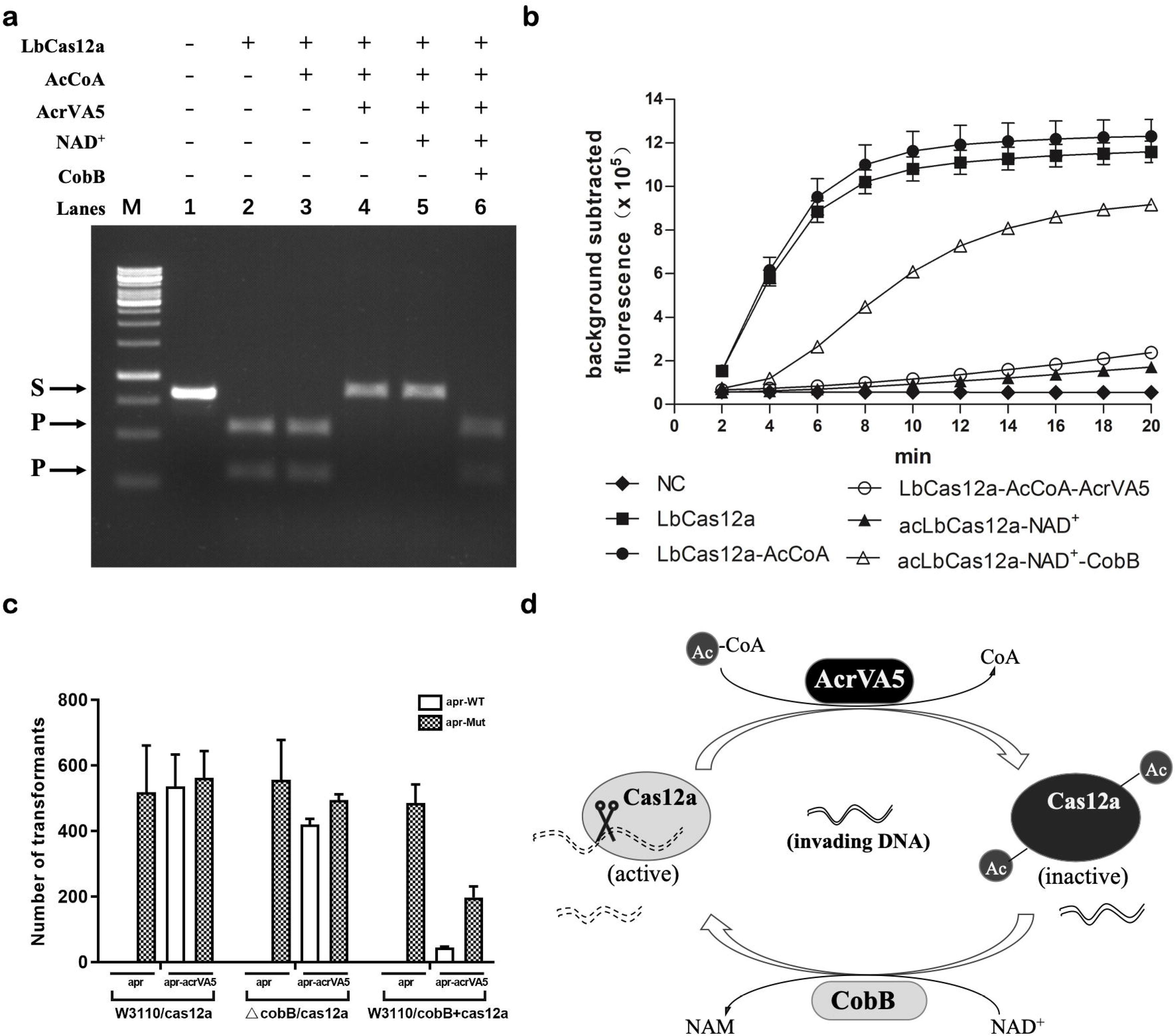
AcrVA5- and CobB-mediated reversible regulation of Cas12a activities through acetylation and deacetylation. **(a)** The LbCas12a *cis*-cleavage experiment with target dsDNA. M, 1-kb DNA ladder (Thermo Fisher Scientific); S, dsDNA substrate; P, Cas12a *cis*-cleaved products. **(b)** The LbCas12a *trans*-cleavage experiment with target dsDNA. Fluorescence signal was collected with a real-time qPCR machine and values were shown with background signal subtracted. NC, the negative control reaction with no target added; LbCas12a, reaction using untreated LbCas12a; LbCas12a-AcCoA, reaction using LbCas12a treated with acetyl-CoA only; LbCas12a-AcCoA-AcrVA5, reaction using LbCas12a treated with AcrVA5 at the presence of acetyl-CoA; acLbCas12a, AcrVA5-acetylated LbCas12a; acLbCas12a-NAD^+^, acetylated LbCas12a treated with NAD^+^ only; acLbCas12a-NAD^+^-CobB, acetylated LbCas12a treated with CobB at the presence of NAD^+^. (c) Analysis of the physiological role of CobB in protecting the host from invading foreign *apr* reporter plasmid. W3110/cas12a, the wild type expressing the LbCas12a/crRNA complex; ΔcobB, the *cobB* null mutant expressing the LbCas12a/crRNA complex; W3110/cobB+cas12a, the wild type expressing the LbCas12a/crRNA complex as well as the CobB. apr-WT, reporter plasmid with the wild type *apr* gene; apr-Mut, reporter plasmid with the mutant *apr* gene that can escape from the cleavage by Cas12a. **(d)** A regulatory model of AcrVA5- and CobB-mediated reversible regulation of Cas12a activities. The acetyltransferase AcrVA5 uses acetyl-CoA to acetylate Cas12a, inactivating Cas12a as well as the host resistance system against MGEs; however, the host possesses the NAD^+^-dependent CobB, which deacetylates and reactivates Cas12a and provides the host a secondary safeguard to the invading nucleic acids.

We further explored the physiological role of CobB in protecting the Cas12a-containing host from the invasion of MGEs that harbor the *acrVA5* gene. We first constructed a CRISPR plasmid that expressed both LbCas12a and a crRNA targeting a specific sequence in the apramycin resistance gene (*apr*) and transformed it into *E. coli* strains with or without a high level of CobB expression (**Figure S7**). Because of the low expression level of the native *cobB* gene in *E. coli*^9^ (**Figure S8**), the wild type W3110 as well as the *cobB*-deleted strains were considered as the *cobB*-negative host, while the *cobB* overexpression strain was the *cobB*-positive host. To prepare a reporter plasmid without the targeting sequence of the Cas12a/crRNA complex, we mutated the Cas12a-targetting DNA sequence in the *apr* gene without changing the amino acid sequence as well as the apramycin resistance (**Figure S9**), obtaining a mutated *apr* gene. Because the nonsense mutation changed both the PAM site and the guide sequence, Cas12a failed to cleave the target sequence both *in vitro* (**Figure S9b**). As a result, the reporter plasmid with mutant *apr* effectively escaped from the host defense system and showed high transformation efficiencies in an *acrVA5*-independent manner (**Figure 1c**), which was therefore employed as a control of the transformation assay.

The reporter plasmid containing the wild type *apr* gene was then transformed into *E. coli* strains expressing Cas12a to mimic the invasion of MGEs, and as expected, in the absence of *acrVA5* gene, no transformants could be obtained in all tested strains independent on the expressional level of CobB, representing the invasion of the foreign plasmid could be completely blocked by the hosts (**Figure 1c**). However, if the reporter plasmid contained both *apr* and *acrVA5*, mimicking MGEs harboring the *acrVA5* cassette, high transformation efficiency of the reporter plasmid was obtained in the *cobB*-negative hosts, which was consistent with the above findings that AcrVA5 effectively inactivated Cas12a. As the overexpression of CobB in the *cobB*-positive host could deacetylate and reactivate the AcrVA5-acetylated Cas12a, the transformation of the reporter plasmid was largely blocked. As expected, the efficiency of the reporter plasmid was still much lower than the control, although overexpression of CobB and AcrVA5 may severely reduce the *E. coli* transformation efficiencies (**Figure 1c**). Based on above results, one may conclude that AcrVA5 and CobB reversibly regulated the Cas12a activities *in vivo* through acetylation and deacetylation, respectively.

As the AcrVA5-mediated acetyl modification of Cas12a has been considered as an effective anti-Cas12a strategy by MGEs,^5, 6^ the CobB-mediated deacetylation reactivation of Cas12a through modification can be taken as both a prevention strategy to the anti-CRISPR elements and a safeguard to the CRISPR defense system, protecting hosts from the invading of the *acrVA5*-containing MGEs (**Figure 1d**). Moreover, this study highlights the potential to create more complexed systems to regulate the Cas12a cleavage activities, which may facilitate both *in vivo* gene editing and *in vitro* target nucleic acid detection.^10^

Besides Cas12a, AcrVA5 may function as a broad-spectrum acetyltransferase and influences cellular metabolic processes. We next performed the *in vitro* western blot analysis through using *E. coli* cellular extracts and purified AcrVA5 and found a large number of proteins could be acetylated by AcrVA5 (**Figure S10**). Similarly, after overexpression of AcrVA5 in *E. coli*, the whole cellular acetylation level was greatly increased, and the acetylated proteins could then be efficiently deacetylated by CobB at the presence of NAD^+^ (**Figure S11**). With the employment of mass spectrometry, we further characterized 2688 sites and 1011 proteins as potential AcrVA5 targets (**Table S2**). Probably due to the small protein size and the reduced steric hindrance in turn, AcrVA5 showed no obvious target sequence preference but favored positively charged amino acids such as lysine, arginine and histidine (**Figure S12**). Considering the extensive targets of AcrVA5, one may imagine that CobB and AcrVA5 compete for control of both the host immune system and beyond. Moreover, the Dsr proteins, which contain the deacetylase domains, was recently found to protect hosts against the invading of dsDNA phages,^11^ and the present study may possibly provide a clue for the unclear mechanisms.

Taken together, one could infer that the wars between invading MGEs and the microbial hosts will never end, and besides acetyl- and deacetyl-modification, there probably exist other kinds of competitive mechanisms subject to future investigation. While just as the Chinese idiom goes, one can believe a vice rises one foot, but virtues rise ten.

## Acknowledgements

We thank Professor Jiaoyu Deng (Wuhan Institute of Virology, CAS) for generously providing the *E. coli* W3110 strain and the mutants. We thank Qiuxiang Cheng (Tolo Biotech.) for her help in data analysis, and Yuangang Ni and Bangtai Luo (Tolo Biotech.) for their help in protein purification, and Jiacheng Wu (Shanghai Tech University) for his assistance in bioinformatics analysis.

This work was supported by grants from the National Natural Science Foundation of China (31922046, 31770057), Sanming Project of Medicine in Shenzhen (SZSM202011017) and the National Key Research and Development Program of China (2018YFA0903700).

