## Supplementary Files for "Reversible regulation of Cas12a activities by AcrVA5-mediated acetylation and CobB-mediated deacetylation"

**Running title:** CobB activates Cas12a by deacetylation

Xiaoman Kang, Lei Yin, Songkuan Zhuang, Tianshuai Hu, Zhile Wu, Guoping Zhao, Yijian Chen, Yong Xu, Jin Wang

### Supplementary information, Materials and Methods

#### Protein purification

Codon-optimized *acrVA5* gene was synthesized by Sangon (Shanghai, China) and cloned into the NcoI/SacI sites of pET28a-TEV expression plasmid. The *E. coli cobB* gene was PCR amplified by primers of cobB-F/R (**Table S3**) and directly cloned into the NcoI/EcoRI sites of pET28a-TEV, obtaining pET28TEV-CobB. To produce recombinant proteins, the protein expression plasmids were transformed into *E. coli* BL21(DE3). Expression of AcrVA5 protein was induced by addition of IPTG, and proteins were purified using the Ni-NTA column and ion exchange chromatography following a similar procedure as previously described.[1] Similarly, EcCobB was produced in the same procedure as we published before,[2] and the Cas12a orthologs were provided by Tolo Biotech (Shanghai, China). Proteins were quantitated using the Bradford method and stored in -80 °C before use.

To express AcrVA5 protein in *E. coli* strains without the T7 RNA polymerase, the T7 promoter of *acrVA5* gene cloned in pET28a-TEV was replaced by the Tac promoter. Briefly, the T7 promoter-containing plasmid was PCR amplified with primers of DS105-tacPf and DS105-tacPr (**Table S3**), the amplicons of which were purified and self-ligated using the EZmax seamless assembly kit (Tolo Biotech), obtaining the plasmid of pTC20004. The Tac-promoted *acrVA5* was expressed in both *E. coli* W3110 and the derived strains such as the *patZ* and *cobB* mutants.

#### AcrVA5-mediated acetylation and CobB-mediated deacetylation

Purified Cas12a was first treated with AcrVA5 in the presence of acetyl-CoA at 25 °C for 10 min in a similar system as previously reported.[1] Briefly, 1 μM Cas12a was treated with 1 μM AcrVA5 with the addition of 2 μM acetyl-CoA in 1× NEB buffer 3.1 (without BSA) in a 10-μL reaction system. Then, acetylated Cas12a was used for either the western blot analysis or *in vitro cis*- and *trans*-cleavage assays. To reactivate acetylated Cas12a by CobB-mediated deacetylation, the AcrVA5-treated Cas12a was directly mixed with 10-μL deacetylation reaction containing purified 4 μM CobB and 0.1 mM $NAD^+$ in 1× NEB buffer 3.1 (without BSA).[3] The deacetylation reaction was carried out at 37 °C for 40 min before the products were used for further analysis.

For cell extracts, both acetylation and deacetylation assays were performed in a similar system as described above except that more proteins were used, e.g. with 27 μg cell extracts, 3 μg AcrVA5 and 20 μg CobB in 30-μL reaction systems.

#### Cas12a *cis*- and *trans*-cleavage assays

For untreated Cas12a, both *cis*- and *trans*-cleavage assays were performed in a similar procedure as we previously described.[2] For assays using AcrVA5- and CobB-treated Cas12a, the reaction systems had a few alterations as described in the following.

The Cas12a *cis*-cleavage was performed at 37 °C for 40 min in a 30-μL system. Briefly, Cas12a was first treated with AcrVA5 in a 10-μL reaction system and then directly added with 10-μL CobB deacetylation system. After ending of deacetylation, a 10-μL *cis*-cleavage mixture, which was prepared by mixing 30 nM target dsDNA, 500 nM crRNA, 10 units RNase inhibitor and 0.5 - 1 μM purified Cas12a in 1× NEB buffer 3.1 without BSA, was directly added into the deacetylated Cas12a system, obtaining a 30-μL *cis*-cleavage system. Reaction was stopped by heating at 85 °C for 5 min and DNA loading buffer was added, followed by gel electrophoresis and imager detection.

Cas12a-mediated *trans*-cleavage assay (i.e. HOLMES assay) was performed at 37 °C in NEB buffer 3.1 without BSA. Similarly, the 10-μL *trans*-cleavage mixture, which is comprised of 0.5 - 1 μM Cas12a, 250 nM crRNA, 10 nM target DNA, 10 units RNase inhibitor and 1 μM HOLMES-P FQ-reporter, was directly added to the 20-μL AcrVA5- and CobB-treated Cas12a, obtaining a 30-μL *trans*-cleavage system. Reaction was monitored in a real-time PCR machine (ABI StepOne Plus) up to 60 min with fluorescence measurements taken every 2 min ($\lambda_{ex}$: 485 nm; $\lambda_{em}$: 535 nm), and the obtained data were presented as background-subtracted with readings taken in the absence of target template.

The dsDNA target was PCR amplified by primers of AMED16s-f and AMED16s-r, purified by gel electrophoresis followed by purification with a gel purification kit, and then quantitated with NanoDrop 2000 (Thermo Fisher Scientific). Single-stranded target DNA of AMED_16S_ssDNA was directly chemically synthesized and used as the template in Cas12a *trans*-cleavage assays. To minimize the influence of the crRNA quality on the assay readouts, crRNA used in this study was chemically synthesized and purified by HPLC from Genscript (Nanjing, China). The crRNA sequence, the HOLMES-P FQ-reporter and the primers used for preparation of the target DNA as well as the plasmid construction were listed in **Table S3**.

**Quantitative analysis of the *cobB* expression level in *E. coli***

*E. coli* strains of the wild-type W3110, Δ*cobB* and the *cobB* overexpression strain (i.e. W3110 harboring plasmid pET28(Tac)-CobB) were first grown in liquid LB medium overnight by shaking at 37 °C, and then inoculated into fresh liquid LB medium with a starting $OD_{600}$ value of 0.1. After about further cultivation for 90 min at 37 °C, the $OD_{600}$ value reached 0.3, and bacterial cells were harvested at 4 °C. Total RNA was extracted using the ZR Fungal/Bacterial RNA MiniPrep kit (Zymo Research), and the trace genomic DNA was removed by treatment with RNase-free DNase I (TaKaRa). Reverse transcription of the total RNA was performed with the PrimeScript II 1$^{st}$ Strand cDNA Synthesis Kit (TaKaRa), after which RT-qPCR was carried out with the SYBR Green qPCR Master Mix (TaKaRa), following the manufacturer's instructions. Three independent biological replicates were performed, and the housekeeping *gapA* gene was used as the internal control. Primer pairs of RT-ECOcobB-F/R, RT-ECOpatZ-F/R and RT-ECOgapA-F/R were used for transcriptional analysis of *cobB*, *patZ* and *gapA*, respectively, using the StepOnePlus Real-Time PCR System (Thermofisher).

**Construction of the plasmids for transformation analysis**

To construct the Cas12a expression plasmid pTC2001, the Cas12a gene was amplified from plasmid pET28a-TEV-LbCas12a with primers of LbCas12a-F/R with FastPfu DNA Polymerase (Tolo Biotech.), and the amplicons were purified and digested with BamHI and SalI before being inserted into the same sites in pQE80L via T4 DNA ligation. Then, the crRNA expression cassette was first prepared by annealing of equal molar crRNA-apr1F/R primers and then introduced into the SalI and HindIII sites of pTC2001, obtaining plasmid pTC20002, which expresses both LbCas12a and a crRNA targeting the apramycin resistance gene (*apr*). The *cobB* gene was excised from plasmid pET28TEV-CobB and cloned into plasmid pET28(Tac)[4], obtaining the plasmid pET28(Tac)-CobB for overexpression of CobB in *E. coli* strains without the T7 RNA polymerase.

The reporter plasmids containing the *apr* gene were constructed by replacing the *ccdB* gene on plasmid p15A-cm-ccdB[5] with the *apr* gene from pBCAm, obtaining plasmid p15A-cm-apr. The oriT cassette from pSET152 was then introduced to the 3'-end of the chloramphenicol resistance gene to generate p15A-apr-oriTf. After that, the TacP-driven *acrVA5* gene was inserted to create the p15A-apr-acrVA5-oriTf.

To mutate the crRNA targeting sequence in the *apr* gene, plasmids of p15A-apr-oriTf and p15A-apr-acrVA5-oriTf were PCR amplified with primers of apr-Mut-F/R, and the amplicons were digested with DpnI (NEB) and then directly transformed into *E. coli* DH10b competent cells, generating target plasmids of p15A-apr(mut)-oriTf and p15A-apr(mut)-acrVA5-oriTf, respectively.

**LbCas12a-mediated *in vitro cis*-cleavage of *apr* gene fragment**

Primers of apr-F/R were employed for amplification of the *apr* gene fragment from plasmids containing the wild and mutant *apr* genes, respectively, obtaining 812-bp products for *in vitro* LbCas12a cleavage.

The 20-μL cleavage system was comprised of 500 nM LbCas12a and 500 nM crRNA, 100 ng wild-type (or mutant) *apr* gene fragment, 10 U RNase inhibitor in 1× reaction buffer. Reaction was performed at 37 °C for 2 h and was stopped by heating at 95 °C for 5 min. Then, DNA loading buffer was added, and the reaction products were analyzed by gel electrophoresis and imager detection. After LbCas12a cleavage, the 812-bp products turned to be two fragments around 300 and 500 bps, respectively.

**Transformation assay**

Plasmid pTC20002 expressing the LbCas12a/crRNA complex was first transformed into the wild-type W3110 or the *cobB* mutant cells, and the transformants were then used to prepare competent cells for subsequent transformation analysis. To mimic the *cobB* high expression environment, plasmids pET28(Tac)-CobB and pTC20002 were co-transformed into W3110.

To test the Cas12a restriction efficiencies, reporter plasmids containing either wild-type *apr* (i.e. p15A-apr-oriTf and p15A-apr-acrVA5-oriTf) or mutant *apr* (i.e. p15A-apr(mut)-oriTf and p15A-apr(mut)-acrVA5-oriTf) were then transformed into above competent cells, using the heat shock method. Specifically, 30 ng reporter plasmids were transformed into 40-μL competent cells, and appropriate amount of the transformed cells were plated on LB plates containing antibiotics of ampicillin (100 μg/mL), apramycin (200 μg/mL) and IPTG (0.3 mM). For example, 1/20 volume of transformants were plated for either the wild type or the *cobB* mutant competent cells, while all transformants of the cobB overexpression strain were plated due to its relatively low transformation efficiency. Noticeably, the expression of Cas12a was induced by addition of IPTG, while apramycin was used for selection of *apr* transformants. Transformants were cultured overnight at 37 °C and colonies were counted for analysis.

**Western-blot assay**

Protein samples were analyzed by 12.5% SDS-PAGE before being transferred to a nitrocellulose membrane (300 mA, 1-2 h). Membrane was first blocked by 1×TBST (20 mM Tris-HCl [pH 7.5], 150 mM NaCl, and 0.1% Tween-20) with 5% non-fat dry milk at room temperature for 30 min and then washed with 1×TBST for 3 times (5 min each time). After that, the membrane was incubated at 4 °C overnight using pan anti-acetyl lysine antibody (PTM Biolab, China) diluted by 1: 1000 in 1×TBST containing 0.05% Tween-20 and 5% non-fat dry milk, followed by wash with 1×TBST for 3 times. Then, the membrane was further incubated at room temperature for 1 h with goat anti-mouse IgG diluted by 1:10000 in 1×TBST containing 5% non-fat dry milk, followed by wash with 1 x TBST for 3 times. Finally, signal was detected by the ECL Western blotting detection system in a luminescent image analyzer (ImageQuant LAS4000 mini, GE Healthcare).

**Quantitative acetylome analysis of *E. coli* expressing AcrVA5 by mass spectrometry**

*E. coli* W3110 harboring pTC20004 was grown at 37 °C in liquid LB medium till $OD_{600}$=0.65. Then, cells were either induced by addition of 0.08 mM IPTG or not induced, and cultured for another 1 h. The $OD_{600}$ values for cells induced and not induced before being harvested were 1.90 and 1.24, respectively.

The mass spectrometry was performed by PTM Biolab. Briefly, harvested *E. coli* cells were stored at -80 °C and then lysed with buffer containing 8 M urea, 1% Protease Inhibitor Mixture III, 3 μM trichostatin A and 50 mM nicotinamide. After centrifugation at 12,000 g for 10 min at 4 °C, the cell debris was removed, and the supernatant was quantitated by a BCA Protein Assay Kit. Equal amounts of protein extracts were precipitated by cold trichloroacetic acid and then washed by cold acetone for three times before being re-dissolved by 8 M urea and 100 mM triethylammonium bicarbonate (TEAB) [pH 8.0].

For trypsin digestion, the protein solution was reduced by 5 mM dithiothreitol at 56 °C for 30 min and alkylated by 11 mM iodoacetamide at room temperature in darkness for 15 min. Then, 200 mM TEAB was used to dilute the protein solution, and trypsin was added at 1:50 (trypsin-to-protein mass ratio) for digestion overnight. Peptides were dissolved in solvent A (0.1% formic acid in 2% acetonitrile) and separated by a nanoElute UPLC system with solvent A (0.1% formic acid in 98% acetonitrile) and solvent B (0.1% formic acid in 100% acetonitrile) at a constant flow rate of 300 nl/min, with the gradient from 6%B to 22%B for the first 42 min, 22%B to 32%B in 3 min, then 32%B to 80%B in 3 min, and finally 80%B for the last 3 min. The peptides were then directly analyzed by NSI-MS with the timsTOF Pro mass spectrometer coupled online to the UPLC, with the electrospray voltage 2.0 kV. The m/z scan range was 100 to 1700 for full scan, and data were collected using the parallel accumulation serial fragmentation (PASEF) model. A data-dependent procedure that alternated between one MS scan followed by 10 PASEF MS scans with 30.0 s dynamic exclusion.

The resulting MS/MS data were processed with the Maxquant search engine (v.1.6.6.0) and tandem mass spectra were searched against Escherichia_coli_strain_W3110_316407 uniprot database concatenated with reverse decoy database. Trypsin/P was specified as cleavage enzyme allowing up to 4 missing cleavages, five modifications per peptide and minimum peptide length was set at 7. The mass tolerance for precursor ions in First search and Main search as well as for the fragment ions was 20 ppm. Carbamidomethyl on Cys was specified as the fixed modification, and oxidation on Met and acetylation modification on the N terminus and the lysine residues were specified as variable modifications. Peptides quantitation was performed using the label free quantitation (LFQ) method and the false discovery rate (FDR) was adjusted to $< 1\%$.

### Supplementary information, Tables

**Table S1. Distribution of the *acrVA5* gene in Mb strains.**

| *M. bovoculi* strains | AcrVA5 | NAD-dependent deacetylase |
|---|---|---|
| 22581 | AKG07142.1 | disrupted |
| 23343 | - | disrupted |
| 28389 | AKG12174.1 | disrupted |
| 33362 | AKG14143.1 | disrupted |
| 57922 | - | AKG15876.2 |
| 58069 | - | AKG19369.1 |
| 58086 | - | AKG17569.1 |

**Table S2. Summary of the quantitative acetylome in *E. coli.***

| | Identified | Quantified | AcrVA5 (induced) / AcrVA5 (uninduced) | |
|---|---|---|---|---|
| | | | up-regulated | down-regulated |
| Sites | 5315 | 2688 | 1241 | 757 |
| Proteins | 1723 | 1101 | 695 | 483 |

**Table S3. Oligonucleotides used in this study.**

| Oligo names | Sequences (5′-3′) |
|---|---|
| AMED16s-f | gtgaactaagccagtagagc |
| AMED16s-r | ctttcgctcctcagcgtcag |
| AMED_16S_ssDNA | cttcttatccaggtaccgtcacttgcgcttcgtccctggc |
| crRNA_AMED_16S_3 | aauuucuacucuuguagaugccagggacgaagcgcaagugac |
| RT-ECOgapA-F | caacgacctgttagacgctgatt |
| RT-ECOgapA-R | acgttcagcggtaacacggatt |
| RT-ECOcobB-F | ctgcgcgagcgtttgcgtc |
| RT-ECOcobB-R | gaaaggtacgaatacctgat |
| RT-ECOpatZ-F | ggccaacacacccaacac |
| RT-ECOpatZ-R | cgatagcggtaattggcgcg |
| cobB-F | cgacgtaggcctttgaattctcaggcaatgcttcccgc |
| cobB-R | attttcagggcgccatggttatgctgtcgcgtcggggtc |
| DS105-tacPf | tacgagccgatgattaattgtcaatttcgcgggatcgagatctcg |
| DS105-tacPr | caattaatcatcggctcgtataatgtgggggaattgtgagcggataac |
| apr-Mut-F | cggaccttggagttgtctctgaTacTttTtggAgATtAccTaatgtaaagcgcagcgcc |
| apr-Mut-R | cgctttacattaggtaatctccaaaaagtatcagagacaac |
| apr-F | atgtcatcagcggtggagtg |
| apr-R | tcatgagctcagccaatcgac |
| crRNA-apr | aauuucuacucuuguagacucugacacauucuggcgccugc |
| LbCas12a-F | gaaatggtggatccatgctgaagaacgtgggcatcg |
| LbCas12a-R | gctttgggtcgactcagtgtttcacgctggtctgagc |
| crRNA-apr1F | tcgacaatttctactcttgtagatgcaggcgccagaatgtgtcagaga |
| crRNA-apr1R | agcttctctgacacattctggcgcctgcatctacaagagtagaaattg |
| HOLMES-P | FAM-tttttt-BHQ1 |

### Supplementary information, Figures

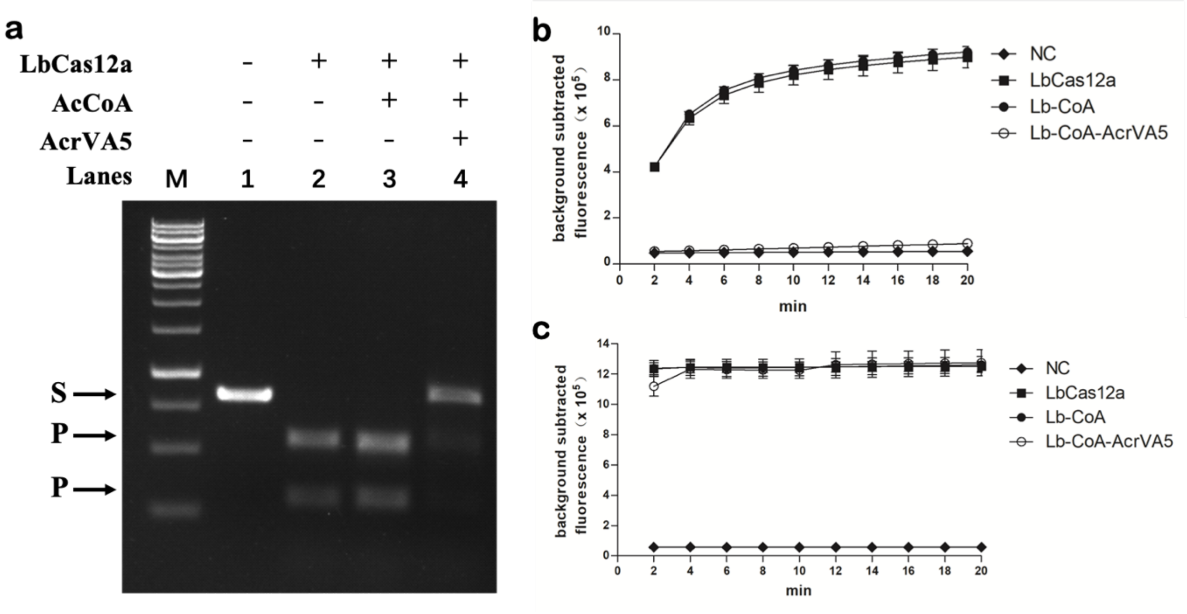


**Fig. S1. The influence of AcrVA5 treatment on LbCas12a *cis*- and *trans*-cleavage activities. (a)** The LbCas12a *cis*-cleavage experiment with target dsDNA. M, 1-kb DNA ladder (Thermo Fisher Scientific); S, dsDNA substrate; P, Cas12a *cis*-cleaved products. **(b)** The LbCas12a *trans*-cleavage experiment with target dsDNA. **(c)** The LbCas12a *trans*-cleavage experiment with target dsDNA. Fluorescence signal was collected with a real-time qPCR machine and values were shown with the background signal subtracted. NC, the negative control reaction with no target added; LbCas12a, reaction using untreated LbCas12a; Lb-CoA, reaction using LbCas12a treated with acetyl-CoA only; Lb-CoA-AcrVA5, reaction using LbCas12a treated with AcrVA5 at the presence of acetyl-CoA.

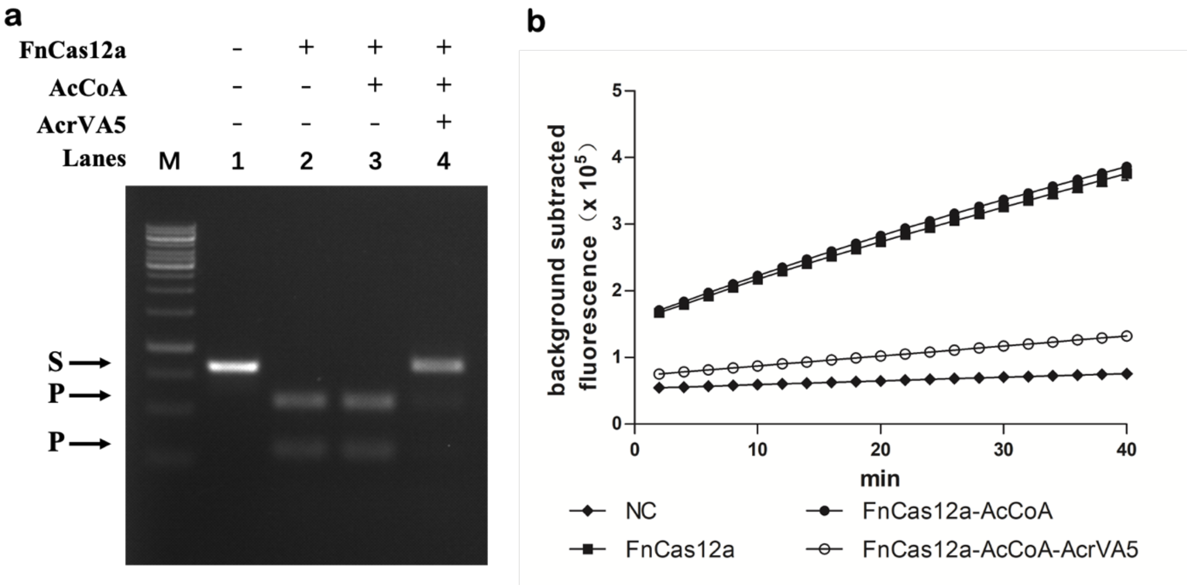


**Fig. S2. The influence of AcrVA5 treatment on FnCas12a *cis*- and *trans*-cleavage activities. (a)** The FnCas12a *cis*-cleavage experiment with target dsDNA. M, 1-kb DNA ladder (Thermo Fisher Scientific); S, dsDNA substrate; P, Cas12a *cis*-cleaved products. **(b)** The FnCas12a *trans*-cleavage experiment with target dsDNA. Fluorescence signal was collected with a real-time qPCR machine and values were shown with the background signal subtracted. NC, the negative control reaction with no target added; FnCas12a, reaction using untreated FnCas12a; FnCas12a-AcCoA, reaction using FnCas12a treated with acetyl-CoA only; FnCas12a-AcCoA-AcrVA5, reaction using FnCas12a treated with AcrVA5 at the presence of acetyl-CoA.

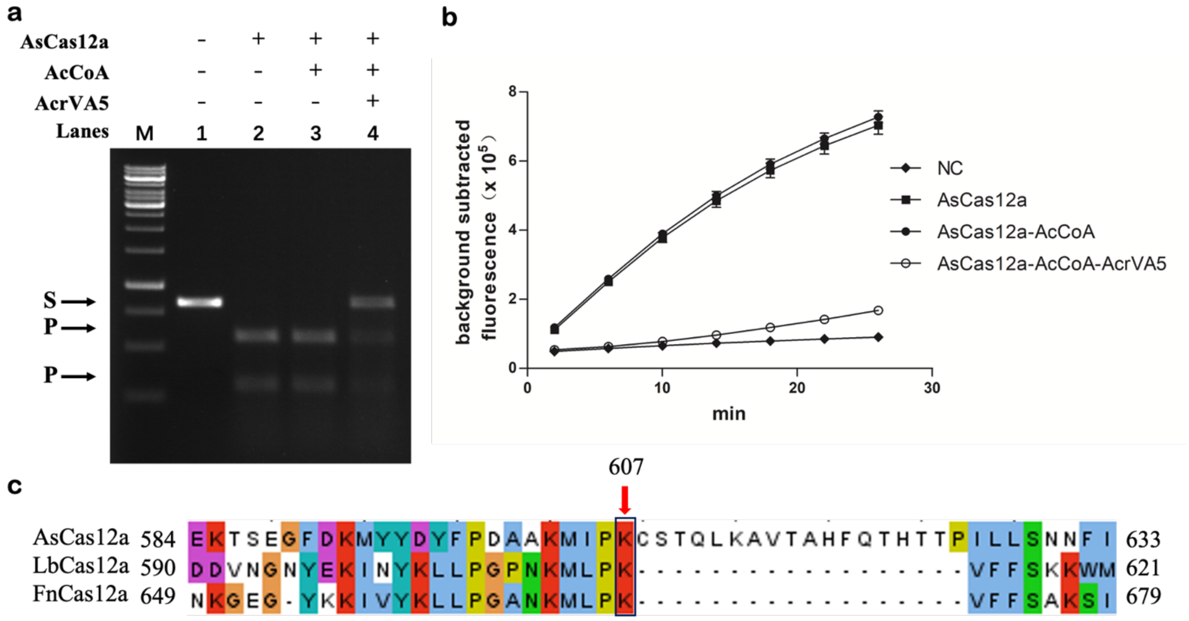


**Fig. S3. The influence of AcrVA5 treatment on AsCas12a *cis*- and *trans*-cleavage activities. (a)** The AsCas12a *cis*-cleavage experiment with target dsDNA. M, 1-kb DNA ladder (Thermo Fisher Scientific); S, dsDNA substrate; P, Cas12a *cis*-cleaved products. **(b)** The AsCas12a *trans*-cleavage experiment with target dsDNA. Fluorescence signal was collected with a real-time qPCR machine and values were shown with the background signal subtracted. NC, the negative control reaction with no target added; AsCas12a, reaction using untreated AsCas12a; AsCas12a-AcCoA, reaction using AsCas12a treated with acetyl-CoA only; AsCas12a-AcCoA-AcrVA5, reaction using AsCas12a treated with AcrVA5 at the presence of acetyl-CoA. **(c)** Sequence alignment analysis of three Cas12a orthologs around the key lysine residue, which was indispensable for PAM recognition and was boxed, and K607 was the residue in AsCas12a.

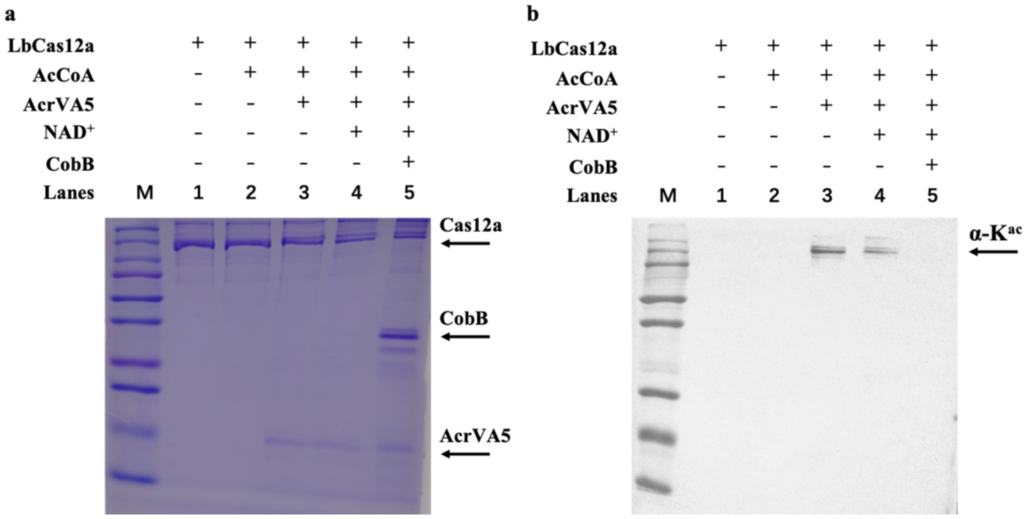


**Figure S4. Western blot analysis of LbCas12a. (a)** The PAGE results of LbCas12a treated with AcrVA5 and CobB. **(b)** The western blot results of proteins in panel **a**. M, the Precision Plus Protein™ standards (Bio-Rad); α-$K^{ac}$, pan anti-acetyl lysine antibody.

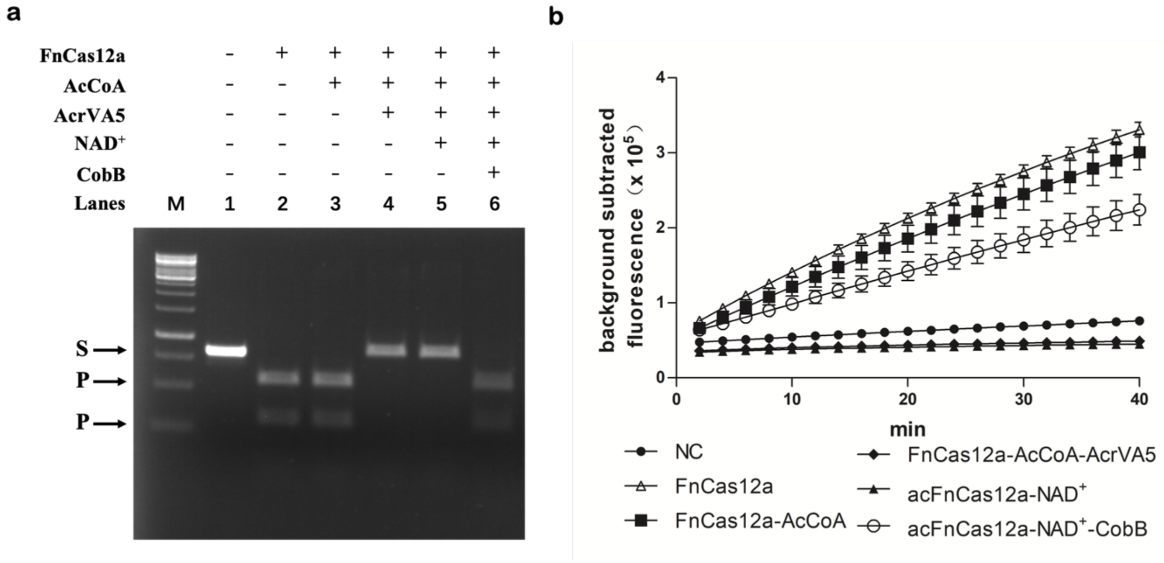


**Fig. S5. The influence of AcrVA5- and CobB-mediated treatment on FnCas12a *cis*- and *trans*-cleavage activities. (a)** The FnCas12a *cis*-cleavage experiment with target dsDNA. M, 1-kb DNA ladder (Thermo Fisher Scientific); S, dsDNA substrate; P, Cas12a *cis*-cleaved products. **(b)** The FnCas12a *trans*-cleavage experiment with target dsDNA. Fluorescence signal was collected with a real-time qPCR machine and values were shown with the background signal subtracted. NC, the negative control reaction with no target added; FnCas12a, reaction using untreated FnCas12a; FnCas12a-AcCoA, reaction using FnCas12a treated with acetyl-CoA only; FnCas12a-AcCoA-AcrVA5, reaction using FnCas12a treated with AcrVA5 at the presence of acetyl-CoA; acFnCas12a, AcrVA5-acetylated FnCas12a; acFnCas12a-$NAD^+$, acetylated FnCas12a treated with $NAD^+$ only; acFnCas12a-$NAD^+$-CobB, acetylated FnCas12a treated with CobB at the presence of $NAD^+$.

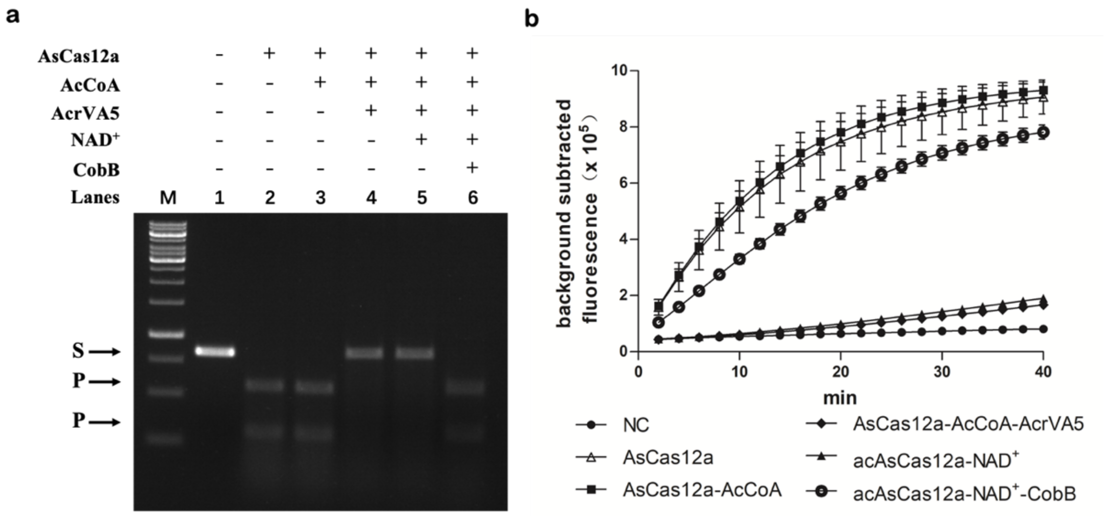


**Fig. S6. The influence of AcrVA5- and CobB-mediated treatment on AsCas12a *cis*- and *trans*-cleavage activities. (a)** The AsCas12a *cis*-cleavage experiment with target dsDNA. M, 1-kb DNA ladder (Thermo Fisher Scientific); S, dsDNA substrate; P, Cas12a *cis*-cleaved products. **(b)** The AsCas12a *trans*-cleavage experiment with target dsDNA. Fluorescence signal was collected with a real-time qPCR machine and values were shown with the background signal subtracted. NC, the negative control reaction with no target added; AsCas12a, reaction using untreated AsCas12a; AsCas12a-AcCoA, reaction using AsCas12a treated with acetyl-CoA only; AsCas12a-AcCoA-AcrVA5, reaction using AsCas12a treated with AcrVA5 at the presence of acetyl-CoA; acAsCas12a, AcrVA5-acetylated AsCas12a; acAsCas12a-NAD$^+$, acetylated AsCas12a treated with NAD$^+$ only; acAsCas12a-NAD$^+$-CobB, acetylated AsCas12a treated with CobB at the presence of NAD$^+$.

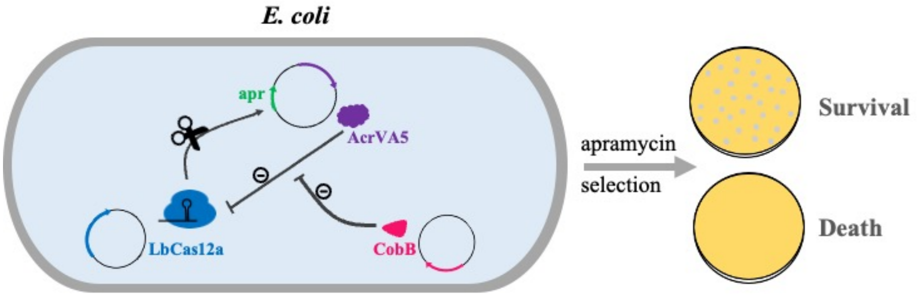


| Strains | *cobB* expression | growth phenotype |
|---|---|---|
| W3110 (WT) | + | +++ |
| *cobB* mutant | - | +++ |
| *cobB* overexpression | +++ | - |

**Fig. S7. Illustration of the Cas12a defending analysis through transformation of the *apr* reporter plasmids.** The LbCas12a/crRNA expression plasmid was first transformed in *E. coli*, which may facilitate the protection of the host from invasion of foreign mobile genetic elements such as the *apr* reporter plasmid in this study. As the reporter plasmid also contained the acrVA5 gene expressing AcrVA5, which acetylated and inactivated Cas12a to allow the transformation of *apr* reporter plasmid and growth of colonies on plates with apramycin selection. While for the strain with a high level of *cobB* expression, the inactivated Cas12a could be reactivated by CobB-mediated deacetylation, protecting the host from the invasion of the foreign plasmid, which would lead to cell death on apramycin selection plates.

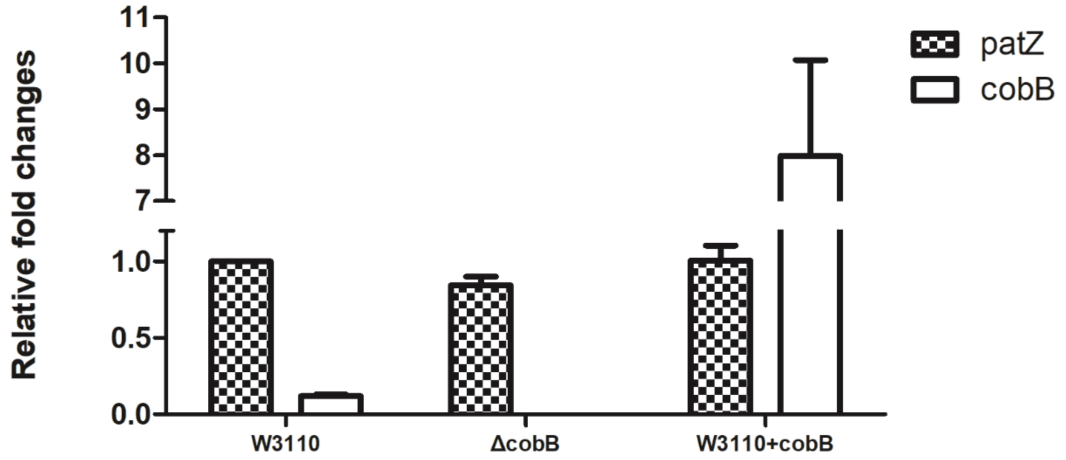


**Fig. S8. Quantitative measurement of the transcriptional level of *cobB* gene and *patZ* gene in *E. coli* strains.** The expression of the housekeeping gene *gapA* was employed as the internal control. W3110, the wild type strain. ΔcobB, the *cobB* mutant. W3110+cobB, the wild type strain W3110 transformed with an overexpress *cobB* (pET28a-tacP-cobB).

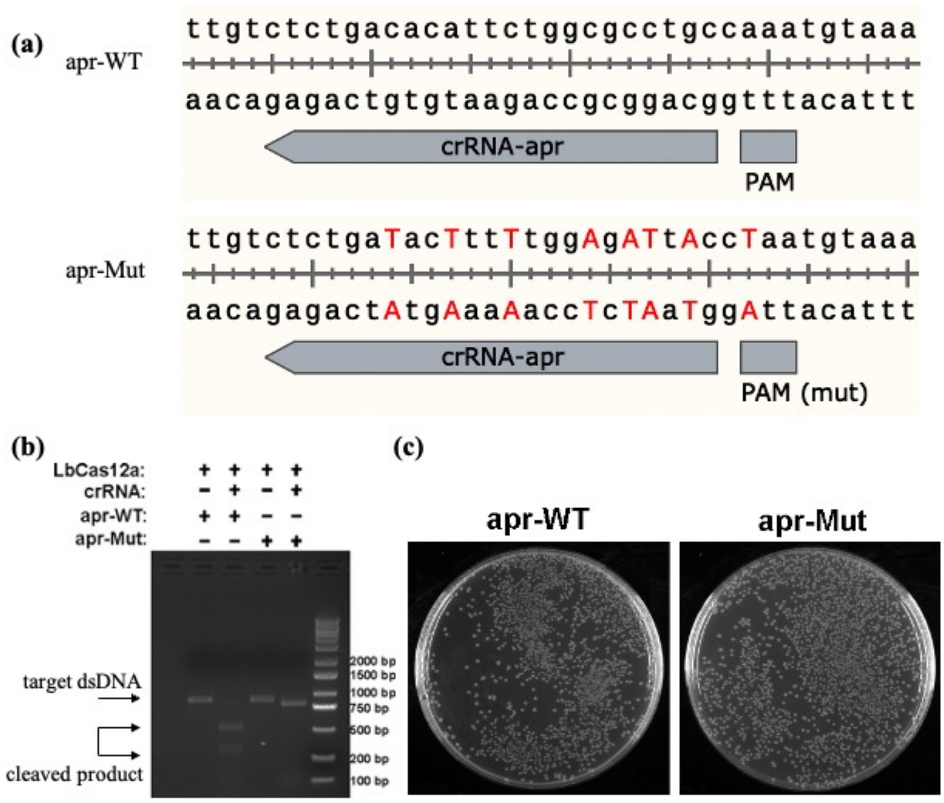


**Fig. S9. Construction of reporter plasmids with mutant *apr* gene. (a)** Illustration of the mutated sequence in *apr* gene, where nonsense mutations were introduced into both the guide sequence and the PAM site of the crRNA-apr sequence. **(b)** Cleavage of the *apr* fragment by LbCas12a. The wild type *apr* fragment instead of the mutated *apr* fragment was successfully cleaved by LbCas12a. **(c)** The apramycin resistance analysis of both wild type and mutated *apr* genes. Both *apr* genes endowed the *E. coli* DH10b hosts with the apramycin resistance, indicating the nonsense mutations did not alter the resistance performances.

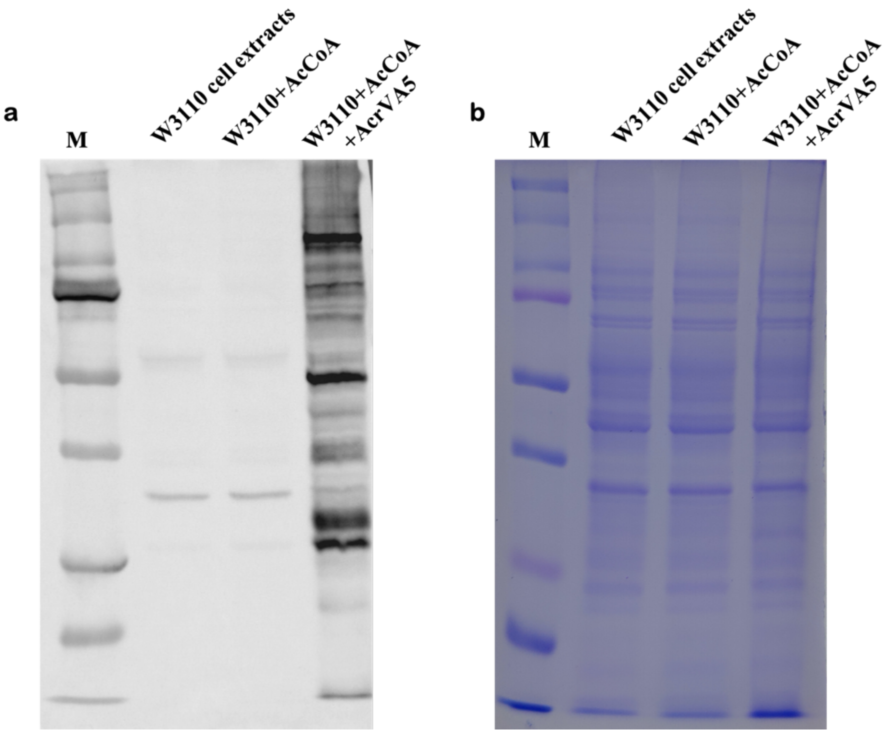


**Fig. S10. AcrVA5-mediated *in vitro* acetylation of *E. coli* W3110 cell extracts. (a)** The western blot results of 27 μg W3110 cell extracts either treated or untreated with AcrVA5, using the pan anti-acetyl lysine antibody. **(b)** The PAGE results stained with Coomassie blue were used as the loading control in panel **a**. M, the Precision Plus Protein™ standards (Bio-Rad). W3110 cell extracts, untreated cell extracts. W3110+AcCoA, W3110 cell extracts treated with acetyl-CoA only. W3110+AcCoA+AcrVA5, W3110 cell extracts treated with AcrVA5 at the presence of acetyl-CoA.

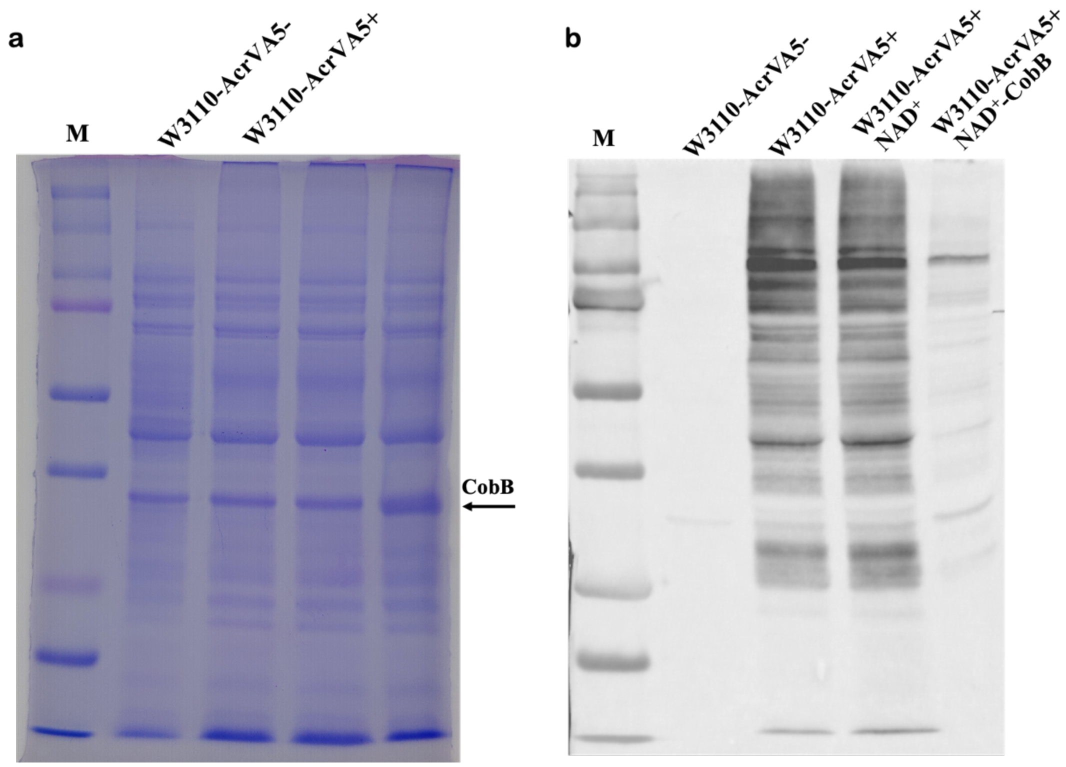


**Fig. S11. CobB-mediated *in vitro* deacetylation of *E. coli* cell extracts. (a)** The PAGE results stained with Coomassie blue, which were used as the loading control of **b**. Thirty microgram cell extracts were loaded and the band of CobB was indicated by a solid arrow. **(b)** The western blot results using the pan anti-acetyl lysine antibody. M, the Precision Plus Protein$^{TM}$ standards (Bio-Rad). W3110-AcrVA5-, W3110 cell extracts. W3110-AcrVA5+, cell extracts of W3110 with the overexpression of AcrVA5. W3110-AcrVA5+NAD$^+$, lane W3110-AcrVA5+ treated with NAD$^+$ only. W3110-AcrVA5+AND$^+$-CobB, lane W3110-AcrVA5+ treated with CobB at the presence of NAD$^+$.

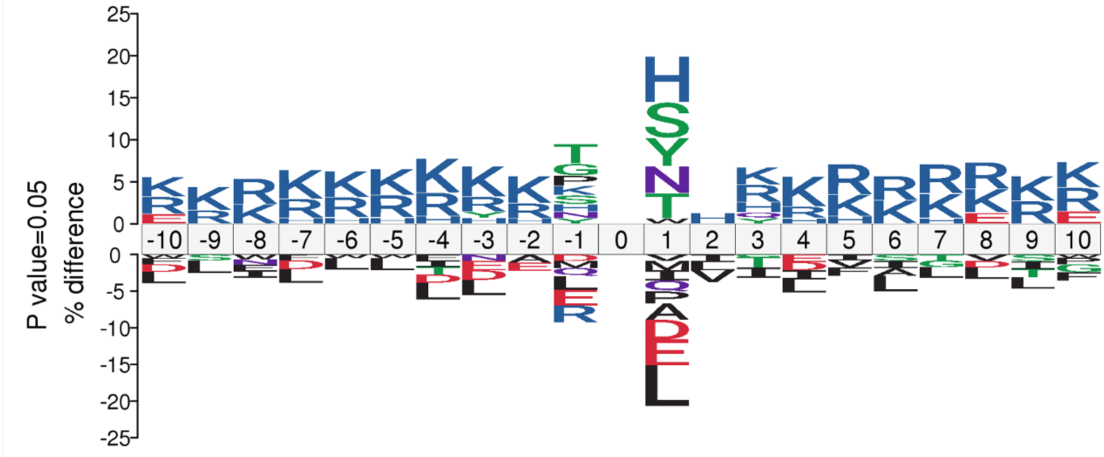


**Figure S12. Analysis of the overrepresented and underrepresented flanking sequences around $K^{ac}$ with the IceLogo software.**
